# Basolateral amygdala neurons await oscillatory recruitment into valence-relevant ensembles

**DOI:** 10.64898/2026.09.15.751884

**Authors:** Kenneth A. Amaya, Yingchu He, Grant L. Weiss, Pantelis Antonoudiou, Jamie L. Maguire

**Author notes:** Corresponding Authors: Kenneth A. Amaya, Jamie L. Maguire.

## Abstract

Impaired valence processing is a core feature of psychiatric illnesses. The ability to rapidly evaluate situations and stimuli and determine whether they have positive or negative implications, termed valence processing, is a highly adaptive brain function that is essential for survival. Despite its importance, we still lack a complete understanding of the neural computations involved. The basolateral amygdala (BLA) plays a critical role in valence processing and valence ensembles have been identified by their anatomical location within the BLA, their projection targets, their genetic identity, or some combination of these factors. In parallel, distinct BLA oscillatory states have been shown to drive divergent valence states. Despite abundant evidence separately supporting these processes, we have failed to reconcile these disparate contributions to valence processing. Here, we demonstrate that subpopulations of BLA principal neurons are recruited in response to specific frequencies of optogenetically-driven oscillations. We provide evidence showing ensemble recruitment is a product of individual neuronal sensitivities to input frequencies that can be driven by interneuron-driven oscillatory states. We also demonstrate that oscillations driven by interneuron stimulations can activate projection-specific populations and reactivate behaviorally-relevant ensembles. Together, these findings reveal a novel neural computational mechanism governing valence processing involving the ability of interneuron-driven oscillatory states to selectively recruit populations of frequency-, projection-, and valence-specific BLA neurons to evoke distinct behavioral outcomes.

## Introduction

Assigning positive or negative valence to experiences is vital to guiding approach or avoidance behaviors and, therefore, survival. Though a broad network of brain regions is involved in valence processing, the basolateral amygdala (BLA) has emerged as a critical hub as it encodes both appetitive and aversive experiences and is densely connected with valence-relevant circuits [1, 2]. Specifically, BLA projection neurons have been shown to divergently guide behavioral outcomes in valenced learning paradigms, where stimulation of BLA projections to the nucleus accumbens (NAc) can promote reward seeking behaviors while stimulation of BLA projections to other regions like the central amygdala, hippocampus, or bed nucleus of the stria terminalis (BNST) can promote fearful responding [3–6]. Valence processing has been studied beyond simply looking at ensembles as defined by their projection target, as ensembles have been identified through their stimulus-evoked physiological responses [7], anatomical location within the BLA [8], and genetic identity [9]. Although the intersection of these factors likely contributes to whether any individual neuron encodes positive or negative valence [10], the physiological mechanisms supporting neuron recruitment to valence-encoding ensembles remain unclear. Recent advancements in areas like the hippocampus and cortex have shown that behaviorally-relevant ensembles may be formed through baseline factors like pre-task activity and the intrinsic excitability of individual neurons [11–13]. In the lateral amygdala, overexpression of transcription factors like CREB in sparse populations has been sufficient to increase fear expression [14], but a gap remains as we do not understand the basis of the baseline differences in neuronal activity which influence their recruitment to or participation in valenced ensembles.

Separately, amygdala physiological activity and microcircuitry have also been tied to valence learning. Oscillations reflect an aggregate of currents from local neuronal populations which can be synchronized by GABAergic interneurons and are involved in the representation, flow, and storage of information [15, 16]. Distinct BLA oscillatory states have been related to valence processing as distinct states are associated with freezing [17–19] and reward seeking [20], with elevated power in low-theta (2-6 Hz) or high-theta (7-12 Hz) frequency bands correlating with the expression or suppression of fear, respectively [18, 19, 21]. Moreover, manipulation of inhibitory interneurons in the BLA, including those expressing somatostatin or parvalbumin, can modulate oscillatory states and, in turn, shape behavioral responses [18, 19, 21–23]. Despite compelling evidence linking oscillations to behavioral states across species, how this process relates to ensemble recruitment and downstream network engagement is unresolved. Specifically, we know little about the interaction between interneurons, neuronal ensembles, and oscillatory states during valence processing. Addressing this gap would further our mechanistic understanding about how regions like the amygdala can serve diverse and oftentimes competing functions while connecting parallel lines research. Here, we provide evidence that links these elements in the BLA to clarify how local circuit dynamics contribute to valence processing through distinct valence- and projection-specific ensemble recruitment to, in turn, influence valence-specific, systems-level network engagement.

## Results

### Valenced learning differentially recruits BLA projection neurons

To investigate whether differently valenced experiences recruited unique subsets of projection neurons in the BLA, we retrogradely labeled BLA neurons projecting to either the nucleus accumbens core (NAc) or the bed nucleus of the stria terminalis (BNST) (Figure 1A) and quantified cFos expression in these projection-specific neurons following fear conditioning or extinction learning (Figure 1B). Behaviorally, both groups of animals (BLA to BNST and BLA to NAc) similarly froze following contextual fear conditioning, shown by a significant effect of Timepoint on Time spent Freezing (F(1, 32) = 131.18; p < 0.001), but not Group (F(1, 32) = 0.0001; p = 0.992), nor interaction between these variables (F(1, 32) = 0.153; p = 0.698) (Figure 1C). For mice that underwent fear conditioning only, the proportion of the cFos ensemble that co-expressed the retrogradely labeled mCherry was greater for BLA-BNST neurons than the BLA-NAc neurons (W = 72; p < 0.001) (Figure 1D), and this was not a product of differing levels of cFos expression among the two groups (W = 41, p = 0.673) (Figure 1E). Animals that underwent extinction learning similarly froze following contextual fear conditioning, shown by a significant effect of Timepoint on Time spent Freezing (F(1, 26) = 86.027; p < 0.001) with no observed effect of Group (F(1, 26) = 0.043, p = 0.837) nor interaction (F(1, 26) = 0.004, p = 0.948) (Figure 1F). Likewise, a linear mixed model revealed that animals in both groups similarly extinguished their fear responding through a significant effect of Session (est: -5.80; 95% CI: -8.50 – (-3.10); p < 0.001) but no effect of Group (est: 5.82; 95% CI: -5.08 – 16.71; p = 0.29) nor interaction (est: -2.10; 95% CI: -5.80 – 1.60; p = 0.26) on Time spent Freezing (Figure 1G). For extinction labeling, the proportion of the cFos ensemble that co-expressed mCherry was greater among the BLA-NAc neurons than BLA-BNST neurons (W = 49; p = 0.0006) (Figure 1H), this was also not a product of varying levels of cFos expression in the BLA overall (W = 28, p = 0.71) (Figure 1I). These data demonstrate that differently valenced behavioral experiences can engage distinct BLA projection populations.

**Figure 1.**
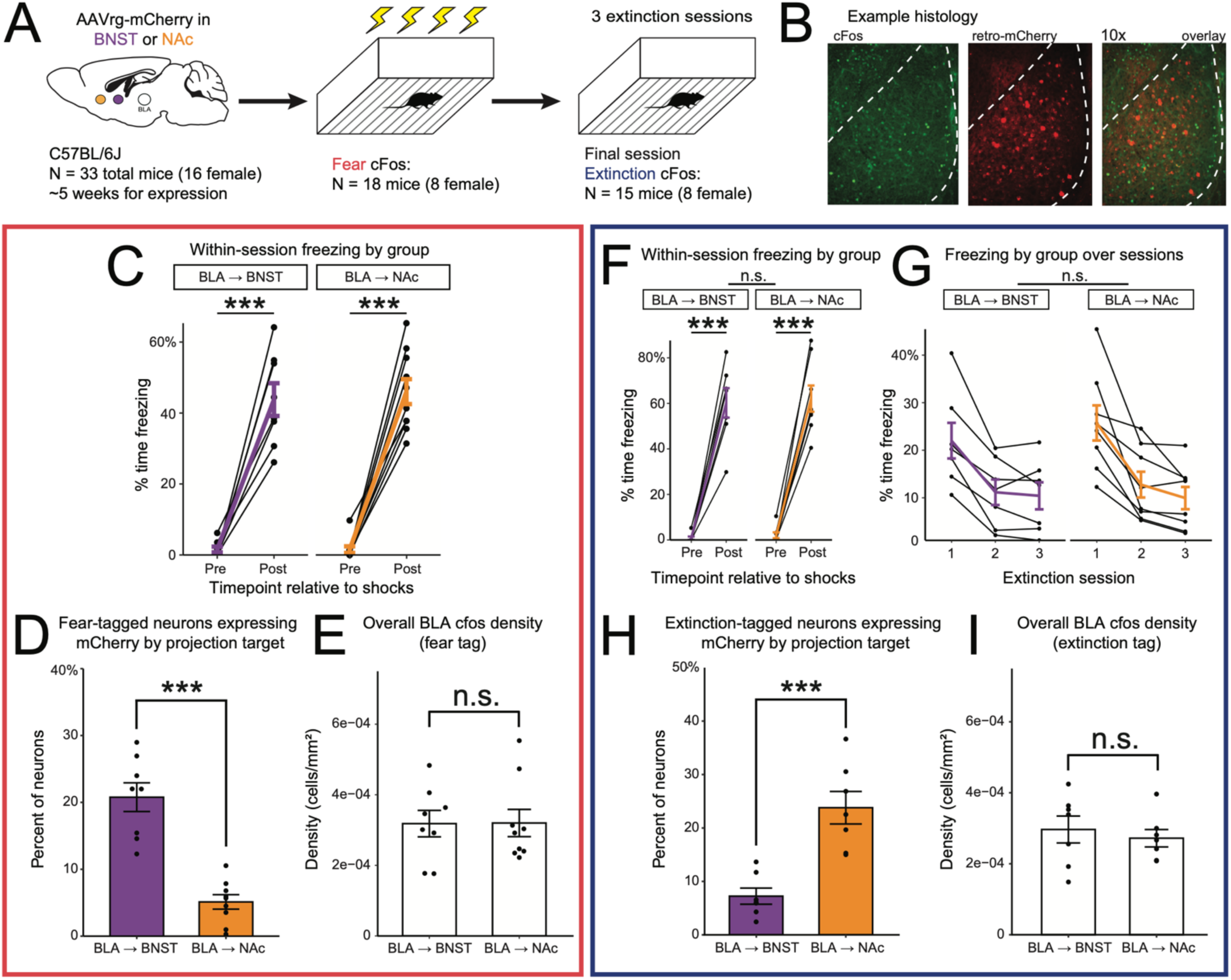
Valenced learning engages distinct BLA projection populations. A) Behavioral timeline showing surgical and behavioral procedures. B) Representative histology showing cFos (left), retrograde-mCherry projections (center), and the overlay of these labels (right) at 10x magnification. C) Proportion of time spent freezing before and after fear conditioning by injection group. D) Fear-recruitment of BLA projection neurons as shown by co-expression of mCherry with cFos. E) cFos density by Group. F) Proportion of time spent freezing before and after fear conditioning by injection group. G) Time spent freezing by extinction session and group. H) Extinction-recruitment of BLA projection neurons as shown by co-expression of mCherry with cFos. I) Mean cFos density by group. n.s., not significant, * p < 0.05, ** p < 0.01, *** p < 0.001.

### Optogenetically-induced oscillatory states reveal BLA neuron tuning preferences

Given that we observed that BLA neurons can be engaged by experience in a pathway-specific manner, we next wanted to understand whether neurons were sensitive to physiological states associated with fear and safety. Fear and safety have been associated with elevated power in the low-theta (2-6 Hz) and high-theta (7-12 Hz) ranges, respectively, and stimulation of parvalbumin-expressing interneurons has been used to artificially entrain the BLA to these network states, bidirectionally biasing behavioral output [18, 19, 21]. Using PV interneuron-induced BLA network entrainment, we investigated whether BLA neuronal populations were recruited during simultaneous *in-vivo* single-photon calcium imaging with optogenetic PV interneuron stimulation at 4 or 8 Hz (Figure 2A). Importantly, we only observed network entrainment in response to 640 nm wavelength stimulations of ChRimson and not to stimulations using 473 nm wavelength light used for calcium imaging (Figure S1). Recordings were conducted in an open arena during a single session that consisted of 10 rhythmic stimulations (five 4 Hz, five 8 Hz stimulations, alternating). From these recordings, we identified a total of 275 cells from 6 mice during recording sessions and found that driving BLA network states through PV interneuron optogenetic stimulation at either 4- or 8-Hz was sufficient to activate unique ensembles. Broadly, cells were classified as either non-responsive or responsive to optogenetically-driven network states, and stimulation-responsive neurons were further parsed into neurons that were indiscriminately responsive to both 4 and 8 Hz stimulations (n = 70; 25.5%) or were preferentially responsive to 4 Hz (n = 35; 12.7%) or 8 Hz (n = 72; 26.2%) (Figure 2B). Responsiveness was not limited to increased activity, as a subset of recorded cells were inhibited by PV stimulations (n = 8; 4.2%) (Figures 2C, 2D). Interestingly, cell responsiveness to PV interneuron stimulation was significantly predicted by pre-stimulation baseline calcium peak events in a logistic regression, such that responsive cells displayed lesser baseline activity than non-responsive cells (Odds Ratio: 0.93, 95% CI: 0.89 – 0.98, p = 0.004). Among responsive cells, baseline activity did not predict whether an individual cell preferred one frequency over another (8 Hz vs 4 Hz: Odds Ratio = 1.07, 95% CI = 0.97–1.17, p = 0.183; Both vs 4 Hz: Odds Ratio = 0.95, 95% CI = 0.85–1.06, p = 0.356) (Figure 2E). Cell preferences were not dynamic within the stimulation and recording period (Figure S2). Mice were naïve to experimental conditions, suggesting that frequency preferences exist at baseline, where neurons may await input to be recruited to state-specific ensembles.

**Figure 2.**
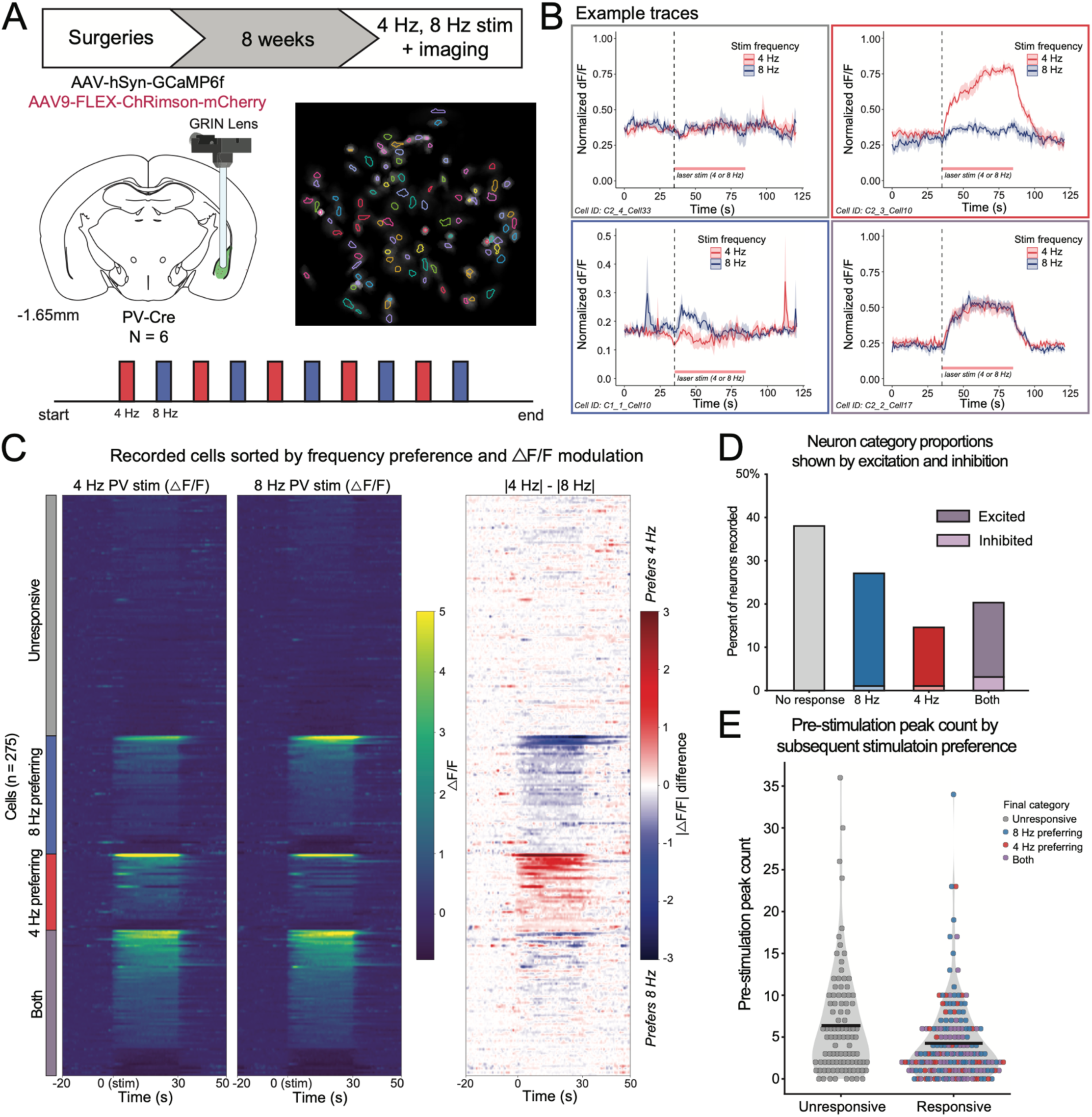
Interneuron frequency stimulations reveal oscillatory preferences among BLA neurons. A) Timeline showing experimental procedures and representative field of view. B) Calcium traces from a non-responsive neuron (top left), 4 Hz-preferring neuron (top right), 8 Hz-preferring neuron (bottom left), and a neuron indiscriminately activated by both frequencies (bottom right). C) Peri-event time heat plot of all neurons, normalized to the pre-stimulation period and averaged over stimulations (4 Hz stimulation, left; 8 Hz stimulation, center). A subtraction of the absolute value of the normalized responses for each cell reveal stimulation preferences (right). D) Proportion of neurons sorted into preference categories. E) Cell responsiveness, but not frequency preference, is predicted by baseline calcium peak events. Black bars represent mean events by condition.

### BLA principal neurons display distinct resonant frequency profiles

The above data provide evidence that BLA projection neurons are differentially recruited by valenced experiences and that artificially-driven oscillatory states can tap into preferential BLA neurons, in general, display preferential activation by unique oscillatory states. Here we assess whether projection- or valence-specific BLA principal neurons are uniquely responsive to frequency-specific input by examining their resonant frequency profiles using whole-cell patch clamp electrophysiological recording. Using this framework, we detail the similarities of the resonant frequency profiles of ensembles defined by projection target, optogenetic/rhythmic recruitment, or experience.

First, we identified BLA projection populations by infusing a retrogradely-expressed fluorescent label into either the BNST or NAc of naïve mice. After allowing for viral expression, acute BLA slices were used for whole-cell patch clamp recordings in current clamp mode (Figure 3A). To understand how labeled, projection-specific neurons respond to oscillatory inputs, we applied a suprathreshold chirp stimulus (increasing in frequency from 1 to 20 Hz across 20 seconds) while recording action potential output from cells in each projection population (Figure 3B). Using a linear mixed model to predict log-transformed normalized spike counts, we observed a significant interaction between Frequency Bin and Projection Target (est: 3.78; 95% CI: 1.10 – 6.46; p = 0.007) and a significant main effect of Frequency Bin (est: -5.66; 95% CI: -7.65 – (-3.67); p < 0.001), but no effect of Projection Target (est: 1.29; 95% CI: -0.61 – 3.18; p = 0.17) (Figure 3B, 3C). This data indicates that BLA projection neurons display projection-specific frequency tuning.

Next, we assessed the electrophysiological properties of neurons that were selectively active during network entrainment at either low or high-theta states. To label these neurons, we optogenetically stimulated interneurons of cFos-TRAP:Ai14 reporter mice at either 4 or 8 Hz in a neutral, novel environment, followed by administration of 4-hydroxytamoxifen (4OHT). Whole-cell patch clamp recordings were performed on Ai14+ frequency-tagged neurons as the chirp stimulus was applied in current clamp mode and action potentials from labeled populations were recorded (Figure 3D). Using a linear mixed model, we observed a significant interaction between Frequency Bin and Tagging Method (est: 4.46; 95% CI: 0.90 – 8.01; p = 0.015), a significant main effect of Frequency Bin (est: -4.16; 95% CI: -6.74 – (-1.59); p = 0.002), but no main effect of Tagging Method (est: 0.22; 95% CI: -2.30 – 2.73; p = 0.86) on log-transformed normalized spike counts during the chirp stimulus (Figure 3E, 3F). Generally, there were no intrinsic membrane differences between the two populations, aside from a significant difference in action potential threshold (4 Hz-tagged: -31.4 mV; 8 Hz-tagged: -36.9 mV) and a significant difference in rheobase (4 Hz-tagged: 67.9 pA; 8 Hz-tagged: 30.8 pA), potentially suggesting that 8-Hz neurons may be more difficult to activate than 4-Hz neurons (Figure S3). These data demonstrate that these frequency-tagged neurons display different sensitivities to inputs, in a manner that is congruent with the frequency used to tag them.

To determine the relevance of these intrinsic frequency signatures to valence processing, we used the same TRAP:Ai14 mouse line to tag BLA ensembles that were activated by either fear or extinction learning. All mice underwent fear conditioning with half receiving 4OHT after conditioning and the remaining half receiving 4OHT after the final of 3 consecutive extinction sessions. Tissue was later collected for whole-cell patch clamp electrophysiology (Figure 3G). Again, we recorded the intrinsic electrophysiological properties of each cell before applying a suprathreshold chirp input stimulus in current clamp mode. Using a linear mixed model, we observed a significant interaction between Frequency Bin and Behavioral Tag (est: -3.75; 95% CI: -7.22 – (-0.27); p = 0.035), but no main effects of Frequency Bin (est: -0.67; 95% CI: -3.13 – 1.79; p = 0.59) nor Behavioral Tag (est: 1.22; 95% CI: -1.24 – 3.67; p = 0.33) on log-transformed normalized spike counts during the chirp stimulus (Figure 3H, 3I). We did not observe intrinsic differences in the electrophysiological properties of the neurons when grouped by tagging method (Figure S4), suggesting that there were no group-based electrophysiological differences other than unique sensitivities to rhythmic inputs.

**Figure 3.**
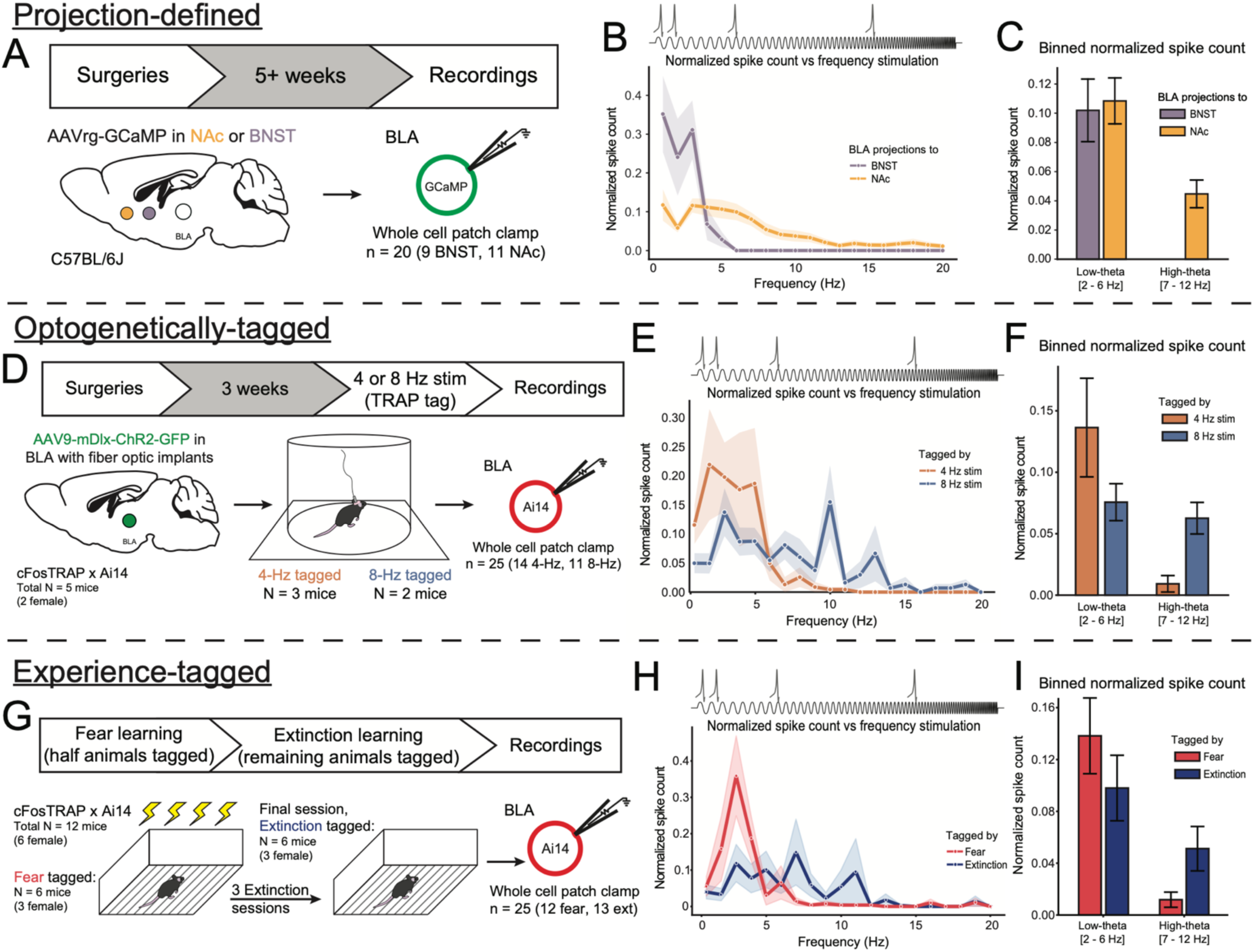
Oscillatory preferences of BLA ensembles are shown through rhythmic stimulation. A) Timeline showing surgical infusion of retrograde-GCaMP followed by ex-vivo whole cell patch clamp electrophysiological recordings. B) Resonance profiles across stimulation frequencies from BLA cells labeled by projection target (BNST-projecting: purple; NAc-projecting: orange). C) Normalized spike count binned by low and high-theta Frequency Bins and shown by Projection Target. D) Timeline showing bilateral surgical infusion of mDlx-ChR2 into the BLA of TRAP:Ai14 mice followed by opto-tagging and subsequent ex-vivo whole cell patch clamp electrophysiological recordings. E) Resonance profiles across stimulation frequencies from BLA cells labeled by optogenetically-induced interneuron recruitment (4 Hz-recruited: red; 8 Hz-recruited: blue). F) Normalized spike count binned by low and high-theta Frequency Bins and Tagging Method. G) Timeline showing behavior-tagging of TRAP:Ai14 mice followed by ex-vivo whole cell patch clamp electrophysiological recordings. H) Resonance profiles across stimulation frequencies from BLA cells labeled by behavioral experience (Fear-tagged: red; Extinction-tagged: blue). I) Normalized spike count binned by low and high-theta Frequency Bins and Behavioral Tag.

### Interneuron-driven network states recruit BLA neurons in behavioral- and pathway-congruent manners

We have shown that animal experience and rhythmic interneuron activity can engage distinct ensembles in the BLA and that frequency-, valence-, and projection-specific ensembles exhibit unique sensitivities to oscillatory inputs. To determine whether interneuron-driven BLA network states can recruit valence-related BLA ensembles, TRAP:Ai14 mice were used to label fear or extinction ensembles prior to undergoing optogenetic interneuron stimulations for subsequent cFos quantification (Figure 4A). Mice were grouped into Congruent and Non-Congruent conditions based on literature linking elevated low-theta power to fear expression while elevated high-theta power is linked to safety (Figure 4B). Mice learned about the threatening context, shown by increases in the proportion of time spent freezing over Timepoint (F(1, 46) = 118.19, p < 0.001), a lack of an effect of Congruency on Freezing (F(1, 46) = 0.0002, p = 0.989), and no interaction between Timepoint and Congruency (F(1, 46) = 0.298, p = 0.86) (Figure 4C). After extinction learning, time spent freezing decreased over sessions similarly between the two groups as a linear mixed model revealed a significant effect of Session on Freezing (est: -8.58; 95% CI: -12.81 – (-4.36); p < 0.001), but no effect of Congruency (est: 14.21; 95% CI: -0.62 – 29.05; p = 0.06), nor was there a significant interaction (est: -4.69; 95% CI: -10.55 – 1.17; p = 0.12) (Figure 4D). After behavioral procedures, BLA interneurons were stimulated at 4 Hz or 8 Hz and tissue was collected for cFos quantification in the BLA (Figure 4E), BNST, and NAc. Mice in the Congruent condition displayed a greater reactivation of BLA (W = 34, p = 0.016) and BNST (W = 26, p = 0.004) behavioral tagged neurons than mice in the Non-Congruent condition, but we did not observe this effect in the NAc (W = 55, p = 0.221) (Figure 4F). We observed coexpression as a product of stimulation frequency and behavioral tag (Figure S5), supporting the notion that Congruence grouping captured a meaningful effect. Together, these data demonstrate that ensembles recruited by behavioral events are re-engaged by interneuron rhythmicity in alignment within a valence-congruent framework.

Given evidence of downstream nodal engagement, we sought to directly assess whether interneuron stimulations can engage distinct projection-specific ensembles. To accomplish this, we retrogradely-labeled BLA projection populations and introduced excitatory opsins for BLA interneuron stimulation (Figure 4G). Following optogenetic interneuron-driven entrainment of the BLA [18–20], tissue was collected for immunohistochemistry labeling of cFos to quantify overlap with BLA-BNST or BLA-NAc neurons (Figure 4H). For analysis, we again grouped animals based on congruency between the projection label and optogenetic stimulations (Figure 4I).

Consistent with our hypothesis, the proportion of projections neurons engaged by interneuron stimulations was greater in the Congruent condition than the Non-Congruent condition (W = 102.5, p = 0.002) (Figure 4J), with a similar pattern observed if analyzing data by projection target and stimulation frequency (Figure S6), suggesting that BLA-BNST neurons are more responsive to 4 Hz network states and BLA-NAc neurons are more responsive to 8 Hz network states. There was no observed difference in the two stimulation frequencies producing more cFos+ neurons (W = 240, p = 0.77) (Figure 4K), nor were there cFos+ neuron differences between the two Congruency conditions (W = 195, p = 0.40) (Figure 4L), suggesting that the difference in activation is best understood through the lens of preferential ensemble recruitment related to the frequency of the network state.

**Figure 4.**
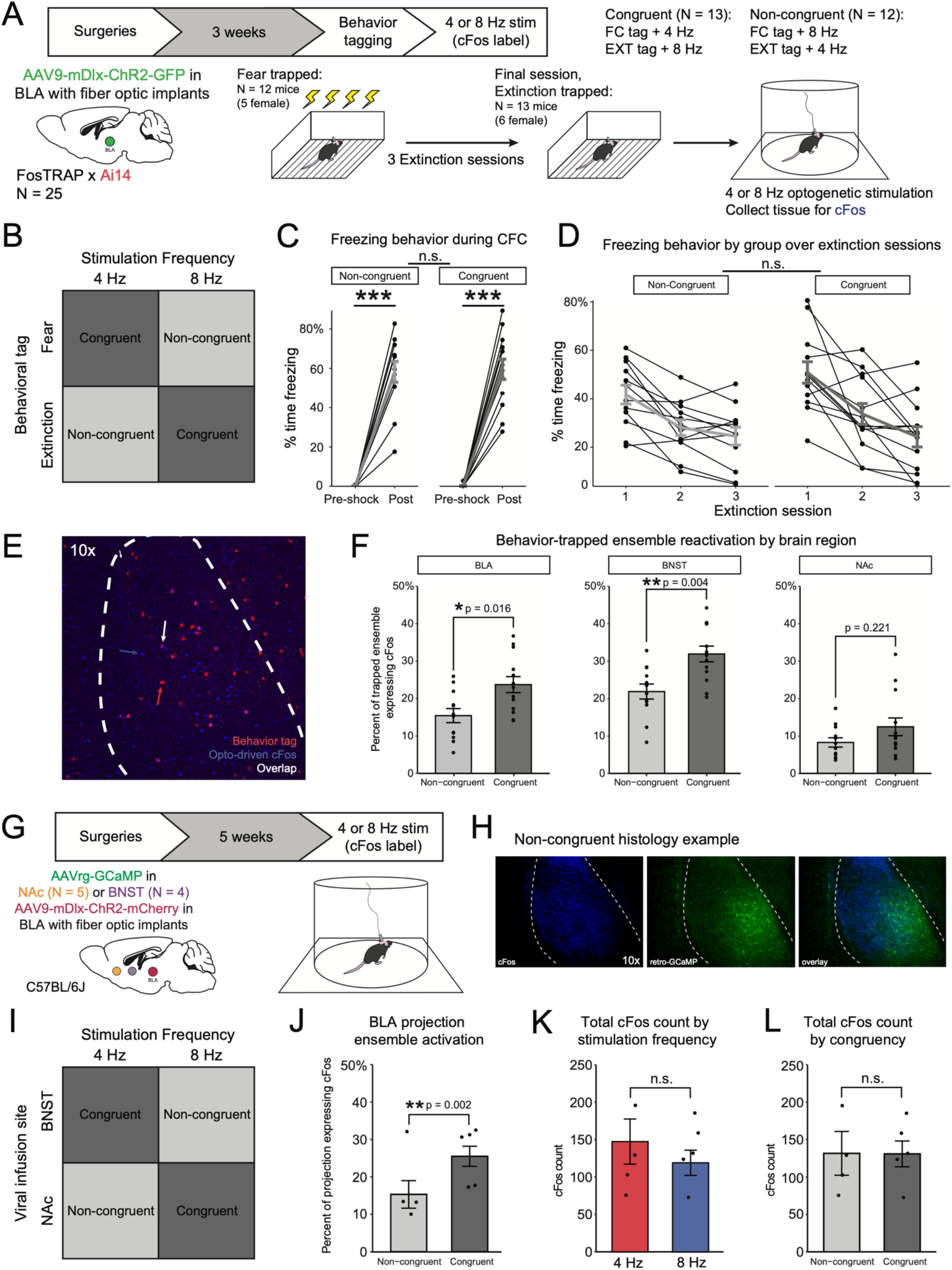
Neuronal ensembles are engaged in congruence with interneuron frequency stimulations. A) Timeline showing surgical, behavioral, and tagging procedures. B) Mice were grouped into conditions for analysis based on behavior and stimulation frequency congruence. Congruent: Fear-tagged/4Hz + Extinction-tagged/8Hz. Non-congruent: Fear-tagged/8Hz + Extinction-tagged/4Hz. C) Freezing behavior before and after fear conditioning by congruency conditions. D) Freezing behavior over extinction sessions by congruency conditions. E) Representative histology at 10x magnification showing an overlay of a behavior-tagged ensemble (red channel, red arrow), optogenetically-induced cFos expression (blue channel, blue arrow), and coexpression (white arrow). F) Ensemble colocalization in the BLA, BNST, and NAc by congruency condition. G) Timeline showing surgical and behavioral procedures. H) Representative histology at 10x magnification showing optogenetically-induced cFos expression (left), labeled BLA projection neurons (center), and the overlay of these labels (right). I) Mice were grouped into conditions for analysis based on projection target and stimulation frequency congruence. Congruent: BNST/4Hz + NAc/8Hz. Non-congruent: BNST/8Hz + NAc/4Hz. J) Ensemble colocalization in the BLA by congruency condition. K) Total cFos count produced by optogenetic stimulation of interneurons. L) Total cFos count by Congruence.

## Discussion

Parallel, yet disparate, lines of research have identified roles for ensembles and oscillatory states in valence processing. While a relationship between physiological network states and behavioral expression of fear or safety has been well described [17, 18, 21], along with previous work highlighting differential involvement of BLA projection populations in this same behavioral paradigm [6, 24], there remained a gap linking network states to behaviorally-relevant ensemble engagement. Using *in-vivo* calcium imaging, whole-cell patch clamp electrophysiology, activity-dependent tagging methods, and fear conditioning and extinction, we found that inhibitory interneurons can engage distinct ensembles in the BLA through their rhythmic activity and that neuronal populations in the BLA are uniquely sensitive to rhythmic inputs. These findings provide evidence for a valence-relevant conceptual framework at the microcircuit level that features the relationship between interneuron and projection neuron. Valence encoding can thus be partially explained through rhythmicity with inhibitory interneurons seated in a prime position as directors of ensemble engagement to support this process.

The idea that ensembles contribute to oscillations is a concept that has been studied in numerous brain regions including the frontal cortex, hippocampus, and ventral striatum [25, 26]. In principle, local spike timing with relation to the excitability of the network is thought to promote neural function, such that spiking during the peak of excitability promotes a winner-take-all model that facilitates nodal communication to engender behavior, often referred to as the communication through coherence theory [27–29]. While prior reports have advanced our understanding of inter-nodal communication, less was known about how intra-nodal ensembles may be selected. We share here that intrinsic biases at the neuronal level may play a key factor in this process. Our group has previously demonstrated that positive and negative experience can reshape the intrinsic electrophysiological properties of BLA neurons related to overall excitability, but not resonant frequency, in a pathway-specific manner [6]. Here, we observed that individual neurons exhibit preferential excitability within specific frequency bands, suggesting that frequency tuning is an inherent feature of BLA principal neurons. This provides a potential mechanism for recruitment of individual neurons into valence-relevant ensembles, whereby oscillatory dynamics may selectively engage neurons based on their intrinsic frequency preferences. Further, the ability of oscillations to engage projection-specific neurons and the established role for oscillations in across brain region communication suggests that this mechanism may also contribute to the distributed engrams associated with valence processing [30]. Consistent with this idea, we observe such recruitment across neurons defined by projection target, behavioral responsiveness, and experimenter-imposed rhythmicity via optogenetic entrainment.

While the current study represents a breakthrough in our understanding of the role of interneurons in valence processing, our knowledge of the role of different interneuron subtypes and microcircuit dynamics is still in its infancy. There are a variety of BLA interneurons, with most identified by markers like parvalbumin (PV), somatostatin (SST), or cholecystokinin (CCK) and there are subgroups even within these interneuron subtypes. We have previously demonstrated that BLA PV interneurons can bidirectionally mediate anxiogenic and anxiolytic network states in the theta range [18, 21], direct beta oscillations for motivated reward seeking [20], and modulate activity in the gamma range via noradrenergic signaling [22, 31]. SST interneurons have been implicated in similar behavioral processes, especially in cue-driven fear expression [23, 32]. Likewise, CCK interneurons have recently been shown to oppose fear learning through modulation of theta power in the BLA [33]. In sum, even of these few interneuron subtypes, there appears to be nontrivial overlap in the ability of GABAergic interneurons to modulate network and behavioral states, especially related to fear learning and expression. Given this, we decided to avoid specific questions about interneuron subtype roles in this study to ask a broader question about how rhythmicity, as generated by interneuron stimulations, can engage ensembles in the BLA. Therefore, most of the work we present here targets interneurons generally through use of the DLX-promoter as part of our viral constructs, though we note that we manipulated BLA PV interneurons in our calcium imaging experiment. This limitation ultimately presents as an opportunity for further inquiry, as it appears that task-specificity may be a deciding factor in which interneuron subtype is predominantly involved in valence processing. We suggest that the principles outlined here, in which interneuron-mediated oscillatory states engage distinct ensembles contributing to valenced behavioral outputs, may inform future work on interneuron subtype contributions to microcircuit dynamics and valence processing in the amygdala.

Despite demonstrating a role for interneurons in orchestrating BLA network states to recruit projection- and valence-specific BLA principal neurons, pressing questions remain regarding the mechanisms mediating the generation of distinct interneuron-driven network states. Specifically, understanding how incoming neuromodulatory signals may influence the activity of interneurons is critical to elucidating how the BLA can transition between network and behavioral states. As a central hub for emotional learning, the BLA integrates diverse afferents, including dopaminergic signaling from the ventral tegmental area (VTA), noradrenergic signaling from the locus coeruleus, and neurotensin from the paraventricular nucleus of the thalamus [22, 31, 34, 35]. Our group has previously reported that VTA dopaminergic transmission to the BLA alters BLA network activity while entraining the frontal cortex during positively-valenced motivated behaviors [34]. Additionally, we have reported that noradrenaline transmission in the BLA shifts oscillatory states to facilitate negative valence processing [22, 31]. And although neurotensin has been convincingly shown as a mediator of valence processing in the BLA, its effects on BLA oscillatory states remains unexplored. Central to how these neuromodulatory systems interact and/or compete with one another to control network states and ensemble recruitment for valenced information routing is the distribution of receptors among cell types. Neurotensin receptors are expressed on principal neurons [35], whereas α1A-adrenergic receptors are expressed heavily on GABAergic interneurons including PV-expressing interneurons, and dopaminergic D1 and D2 receptors are found on both principal neurons and fast-spiking interneurons [36]. Together, the relative distribution of these receptors coupled with the temporal dynamics of neuromodulator release may dictate how these inputs sculpt oscillatory states, recruit valenced ensembles, and ultimately govern downstream network engagement.

Valence processing has been recognized as a major domain in the Research Domain Criteria framework to further classify mental disorders [37]. We have previously highlighted that the neural circuitry involved in valence processing is remarkably similar to aberrant neural circuits in psychiatric conditions, with the amygdala seated centrally among both [38]. In preclinical research, fear/threat conditioning and extinction learning has been used as a model for disorders like post-traumatic stress disorder, given that mice will readily learn about a threat in the environment but will readily reduce their threat responding as extinction progresses. Presently, we observe that distinct ensembles, identified across multiple definitions, are involved in these learning processes, with interneurons excitingly positioned to contact and reactivate ensembles in an oscillatory state specific manner. Given that oscillations are one of the most widely observed types of neural activity dynamics across species [39], characterizing oscillatory states related to valence processing has strong translational implications for understanding rigid post-traumatic stress responses and for neuromodulatory therapeutic opportunities. Mechanistic inquiries into the generation and regulation of such fearful neural states, including characterizing the roles of interneurons in cortical and subcortical limbic networks, could provide an avenue for therapeutic advancement through the development and application of compounds or interventions that sway networks back to a healthy balance [21].

Overall, the findings reported here are consistent with prior work that identified BLA neurons as active in distinct behaviorally-relevant oscillatory states [19]. Using this framework, we detail that rhythmic interneuron activation can engage distinct BLA ensembles tied to valence learning as defined by their projection target, sensitivity to rhythmicity, or encoding of behavioral experience. Our work offers insights into the relationships between ensembles, oscillations, and behavioral output, with interneurons seated centrally in mediating their interactions, ultimately offering a stronger understanding of the ensemble-oscillation relationship that supports learning and behavior.

## Methods

### Subjects

Adult (greater than 10 weeks old) male and female C57BL6/J mice (C57; #000664; N = 51), B6.129P2-Pvalbtm1(cre)Arbr/J (PV-Cre; #017320; N = 6), Fos^2A-iCreER^(TRAP2) (TRAP; #030323), and B6.Cg-Gt(ROSA)26Sor^tm14(CAG-tdTomato)Hze^ /J (Ai14; #007914) were acquired from The Jackson Laboratories. TRAP and Ai14 mice were bred in-house to produce TRAP:Ai14 activity-dependent reporter mice used in these experiments (N = 42). All animals were housed at Tufts University School of Medicine in an AAALAC-approved temperature and humidity-controlled facility with a 12-hour light/dark cycle with lights on at 7 AM, with food and water provided *ad libitum*. All procedures were approved by Tufts University’s Institutional Animal Care and Use Committee (IACUC).

### Surgical procedures and viral vectors

Mice that underwent surgical procedures were anesthetized via intraperitoneal injection of a ketamine/xylazine cocktail (100mg/kg ketamine and 10 mg/kg xylazine) and received subcutaneous sustained release buprenorphine (0.5 mg/kg) prior to surgery. The basolateral amygdala (BLA) was stereotaxically targeted using these coordinates relative to bregma (AP: -1.50; ML: ± 3.0; DV: -4.5). Additionally, the following coordinates were used to target nucleus accumbens core (AP: +1.7; ML: ± 0.75; DV: -4.0) and the bed nucleus of the stria terminalis (AP: +0.02; ML: ± 0.5; DV: -4.0).

For *in-vivo* calcium imaging and optogenetic stimulations, mice were unilaterally injected with 500 nL of a 1:1 viral cocktail containing AAV-DIO-ChRimson-tdTomato and AAV9-hSyn-GCaMP6f (Addgene #105448 and #100837) into the BLA using a 33-gauge Hamilton syringe at an infusion rate of 100 nL/min. After allowing 10 minutes for viral diffusion, an Inscopix lens affixed to a baseplate (Inscopix, 1050-004413) was placed above the BLA. Mice were allowed to recover for at least 6 weeks prior to experimentation to allow for optimal viral expression and *in-vivo* imaging and optogenetic manipulation.

For optogenetically induced ensemble tagging experiments, mice were bilaterally injected with 250 nL of AAV9-DLX-ChR2-mCherry (Addgene #83898) or AAV9-DLX-ChR2-EYFP (Harvard viral core) into the BLA using a pulled glass pipette with a 20 μm diameter opening at an infusion rate of 100 nL/min. Following each infusion, the glass pipette was allowed to rest for an additional 10 minutes before removing from the brain. Two anchor screws were placed into the skull and fiber optics (200 μm, 0.22 NA; ThorLabs) were bilaterally positioned over the BLA and cemented in place. Animals were allowed at least 3 weeks to recover and for viral expression.

For retrograde labeling of BLA projection populations, mice received 250 nL a AAVrg-hSyn-GCaMP6f (Addgene #51085) or AAVrg-hSyn-mCherry (Addgene #114472) into either the NAc or BNST, depending on the experiment. Differing viral labels were used to accommodate other viral constructs and avoid label overlap. For example, when assessing overlap between optogenetically-recruited ensembles and projection populations, given the viral construct we had at the time for targeting interneurons (AAV9-DLX-ChR2-mCherry), we used a GFP-related projection label in AAVrg-hSyn-GCaMP6f. Animals that received a retrograde label for projection population identification were allowed at least 5 weeks for viral expression.

### In-vivo calcium imaging

On the day of recordings, each animal was removed from their home cage, had a miniature integrated microscope system (nVoke HD 2.0; Inscopix) attached to their baseplate, and was given 10 minutes to habituate to the testing cage (identical to the home cage with fresh bedding and nesting). Images were acquired using the data acquisition software (version 2.0.0; Inscopix) at 20 frames per second, 20% of LED power, and a gain kept between 2 and 5, varying based on the clarity of the field of view.

Calcium imaging recording sessions were 10-15 minutes in length and included 10 total optogenetic stimulation periods that each lasted 30 seconds in duration. Stimulations were delivered Acquired imaging data were down-sampled (1/2 spatial binning), preprocessed, motion corrected, and then calcium transients of individual neurons were extracted via PCA-ICA using Inscopix Data Processing Software (version 1.9.1; Inscopix). All extracted traces were manually checked resulting in traces from multiple cells or non-cellular signals being excluded. The fluorescent trace data was then subjected to a series of custom-built scripts (implementing SciPy’s peak-finding algorithms in Python) which automated the detection of calcium transient events, generating a structure which detailed the peak amplitude (height), area under the curve (AUC), width, peak-to-peak prominence, rise time, and decay time for each trace’s events within a given recording.

### Ensemble tagging methods

For TRAP:Ai14 animals that were involved in activity-dependent ensemble tagging, 4-hydroxytamoxifen (10 mg/mL; Sigma Aldrich #H6278) was administered (50 mg/kg) immediately after conclusion of optogenetic stimulations or after behavior concluded. Animals were returned to their homecage and were undisturbed for at least the following 3 days to reduce possible inadvertent ensemble labeling. For mice that underwent immunohistochemistry labeling of cFos+ neurons, those animals were sacrificed 90 minutes after the completion of optogenetic stimulation or behavior.

### Ex-vivo electrophysiology

Mice were deeply anesthetized with isoflurane, decapitated and the brain was rapidly removed and placed immediately in ice-cold, oxygenated normal artificial cerebrospinal fluid [nACSF; containing, in mM, 126 NaCl, 26 NaHCO_3_, 1.25 NAH_2_PO_4_, 2.5 KCl, 2 CaCl_2_, 2MgCl_2_, and 10 dextrose (300-310 mOsm)] with 3 mM kynurenic acid and bubbled with 95% O_2_ – 5% CO_2_. Coronal BLA sections (350 μm in thickness) were prepared using a Leica VT1000S vibratome and allowed to recover for at least 1 hour prior to recording. Electrophysiological recordings were performed in a recording chamber maintained at 33°C (in-line heater; Warner Instruments) and perfused using nACSF (external solution) at a high flow rate (∼4 mL/min) throughout the experiment. Borosilicate glass electrodes were pulled with a resistance of ∼3 – 5 MΩ (DMZ Universal Puller). The intracellular recording solution contained (in mM) 130 potassium gluconate, 10 KCl, 4 NaCl, 10 HEPES, 0.1 EGTA, 2 Mg-ATP, and 0.3 Na-GTP (pH: 7.25; 280 – 290 mOsm). Intrinsic electrophysiological properties were measured in visually identified BLA neurons based on morphology and expression of fluorescent tags (GFP or tdTomato). Series resistance and whole-cell capacitance were continually monitored and compensated throughout the course of the experiment. Recordings were excluded from data analysis if series resistance increased by >20% or if patched cells were deemed to be interneuron-like, as determined by their interspike intervals and waveform properties. Data acquisition was carried out using an Axopatch 200B (Axon Instruments) and PowerLab hardware and software (ADInstruments) at 10 kHz sampling. To characterize cell frequency sensitivities, physiological responses (spikes) were measured in response to the application of a suprathreshold chirp stimulus (1 – 20 Hz) over the course of 20 seconds. Data analysis was performed using Python analysis scripts developed in-house. Action potentials were detected using SciPy’s *find_peaks* function (prominence = 50 mV, wlen = 100 ms, distance = 1 ms). Total cells recorded: n = 70.

### Fear conditioning and extinction

Mice were subjected to a 10-minute contextual fear conditioning session in a square chamber (120 cm x 100 cm x 120 cm) with a grid floor (Coulbourn Instruments; H10 – 11RTC). Each chamber was outfitted with an overhead digital camera connected to a computer that also operated the events in the chambers via ActiMetrics FreezeFrame software (ver. 5.104). During contextual fear conditioning after 4 minutes, 4 shocks (2s, 0.70 mA) were delivered with an inter-shock interval of 90 seconds. Following contextual fear conditioning, mice underwent 3 daily sessions of extinction learning (15 minutes in duration) in the same chambers and context as fear conditioning took place.

Animal movement was recorded for quantification of freezing behavior using FreezeFrame software. For each mouse, the motion threshold was manually calibrated by visual inspection of the video in parallel with automated scoring to ensure accurate freezing classification. During contextual fear conditioning, freezing was binned into 60-second pre- and post-shock time windows. Freezing during extinction is presented for the entire 15-minute extinction session over days. Data are presented as mean freezing duration in seconds and the standard error of the mean (SEM).

### Immunohistochemistry

For quantification of cFos+ neurons, animals were sacrificed 90 minutes after the completion of optogenetic stimulation or behavior. Brains were extracted and post-fixed in 4% PFA for 24 hours before being transferred to 1x PBS. Coronal sections (50 μm) were collected using a Leica VT1000S vibratome and stored in 24-well plates at 4 °C. Free floating sections were incubated in blocking solution (phosphate buffered saline, 0.2% Triton-X, and 5% Normal Goat Serum) on a shaker at room temperature for 1 hour. Sections were then incubated in primary antibody solution (blocking solution, 1:1000 rabbit polyclonal anti-cFos [Synaptic Systems #226008]) on a shaker at 4 °C for 48 hours.

Sections were then rinsed 3 times (10 minutes per) in 1x PBS and transferred to secondary antibody solution (blocking solution, 1:200 Alexa goat anti-rabbit 488 [Invitrogen #A11008]) covered in foil on a shaker at room temperature for 2 hours. Sections were then rinsed 3 more times in 1x PBS, mounted onto slides, and cover-slipped with a hard-set mounting medium with DAPI (Vectashield #H-1500-10).

For one experiment, we substituted out the 488-secondary for a 405-secondary (Invitrogen #A48254) at the same concentration and for the same incubation duration. Sections were similarly rinsed and mounted, but were cover slipped with a clear hard-set mounting medium that did not include DAPI (ProLong Glass, Invitrogen #P36984).

### Microscopy and quantification

Images of the basolateral amygdala, nucleus accumbens, and bed nucleus of the stria terminalis were acquired at 10x magnification using a wide-field epifluorescence microscope (Leica THUNDER Imager Tissue). Acquisition settings were optimized for each brain region and were identical across groups. Images were collected bilaterally from each brain region, coming from a minimum of 2 BLA sections, 2 NAc sections, and 2 BNST sections. A small number of images were excluded for tissue quality reasons. CellProfiler (ver 4.2.6, Broad Institute, Cambridge, MA) was used to identify and quantify cFos positive neurons in each brain region. After applying experimenter-drawn masks to restrict quantification to regions of interest, cells were identified with diameter constraints (4-15 pixel units in BLA; 3-9 pixel units in NAc; 4-15 pixel units in BNST), using built-in thresholding methods (Robust Background). This method was applied to all images and produced counts of cFos+ neurons from the left and right hemispheres of the brain region. In animals where we were interested in looking at ensemble overlaps between labeling techniques (e.g., fear-activated ensemble overlap with optogenetically-activated ensemble), we used a different filter cube to quantify expression of the viral or endogenous label. After counts and overlaps were quantified, we averaged values from each section to produce a single overlap percentage for each animal. Those animal-level values were used to quantify the mean and SEM shown in each Figure.

### Statistical analyses

Statistical analyses were conducted in R (R Core Team 2016; “stats”, “lme4”, “lmerTest”, “car”) or Python (“statsmodels”). Figures were generated using R (“ggplot2”) or python (“seaborn”) and stylized in Adobe Illustrator. Unpaired Student’s *t*-tests or Wilcoxon rank-sum tests were used to compare means between two experimental groups and one-way ANOVAs were used to compare means among multiple factors. Where appropriate and noted, linear mixed models were employed to account for individual mouse differences.

For calcium imaging, stimulus-evoked response preferences were determined by averaging ΔF/F during the stimulation window (30 sec in duration), yielding a single response value per neuron, frequency, and trial. Neuronal responsiveness was defined as exceeding a threshold of 0.25 in the absolute mean ΔF/F response in at least one stimulus condition. Neurons were further classified based on their relative responses to the two frequency stimulations: 4 Hz-preferring neurons exhibited responses to 4 Hz that were at least 1.25 times larger than responses to 8 Hz, 8 Hz-preferring neurons exhibited the reciprocal relationship, and neurons without a clear dominance were classified as “Both,” whereas neurons below threshold in both conditions were considered unresponsive.

Following classification, pre-stimulation peaks were used to determine whether baseline activity was predictive of eventual neuron classification. Binary logistic regression was used to test for predictiveness of baseline activity on responsiveness. Models were estimated using maximum likelihood under a binomial error distribution with a logit link function. Statistical significance of model coefficients were assessed using Wald z-tests as implemented in the model summary output. Reported effects were determined as log-odds and exponentiated to yield odds ratios for interpretability.

Data in figures are shown as the mean and standard error of the mean (SEM). p-values of < 0.05 were considered statistically significant and denoted as p ≥ .05 = n.s., *p* < .05 = *, *p* < .01 = **, and *p* < .001 = ***.

## Acknowledgements

This work was supported by the National Institute of Mental Health (JLM: R01 MH128235; KAA: K00 MH130162).

## Contributions

Conceptualization: KAA and JLM

Methodology: KAA and JLM

Data collection: KAA, YH, GLW, and JLM

Data analysis: KAA, GLW, and PAA

Funding acquisition: KAA and JLM

Writing and revising: KAA and JLM

## Supplemental Figures

**Supplemental Figure S1.**
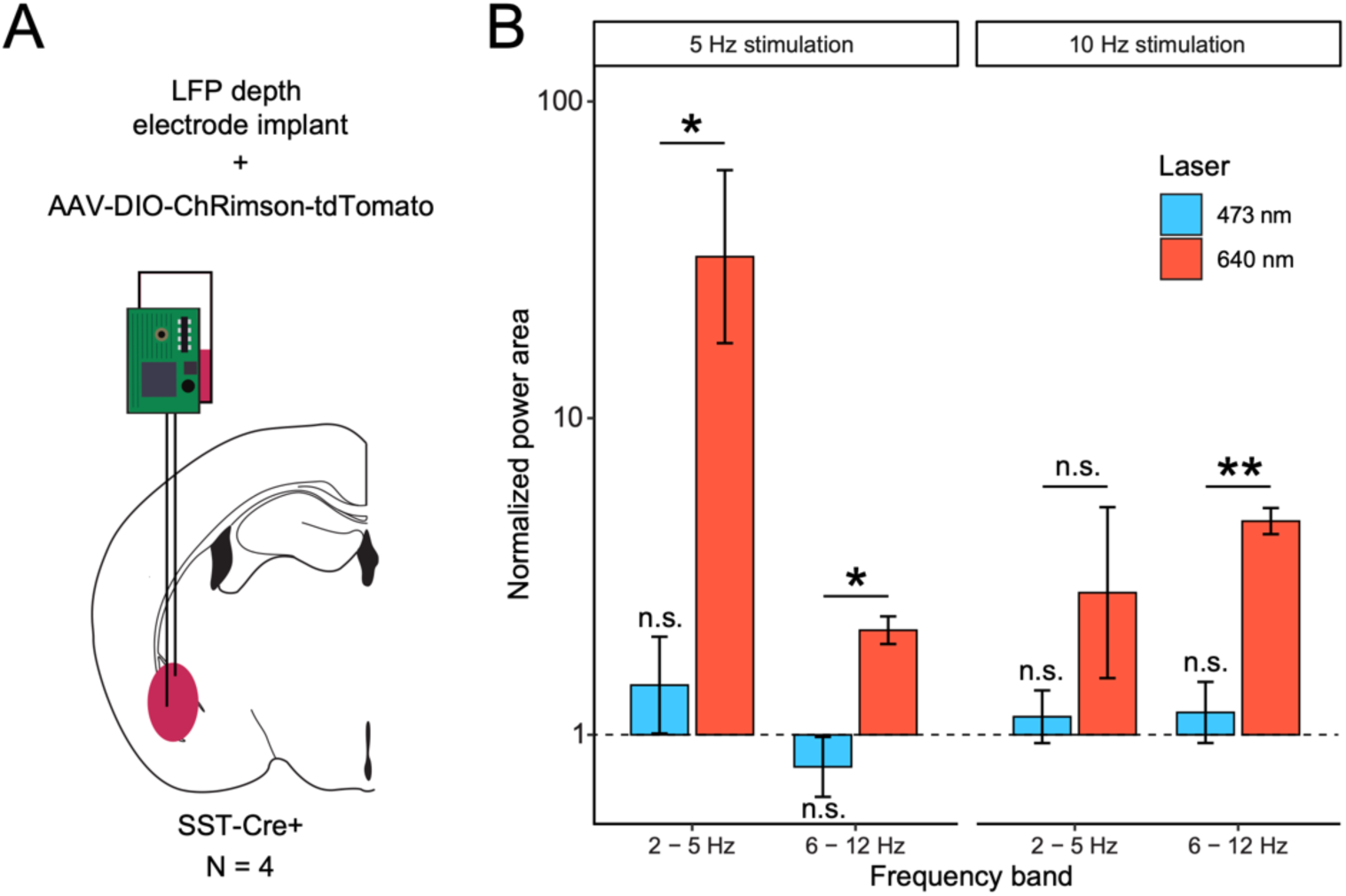
Administration of blue light (473 nm) to ChRimson-expressing interneurons does not alter network states. A) Depiction of procedures where Cre-dependent ChRimson was infused into the BLA of SST-Cre+ mice (N = 4) followed by implantation of a depth electrode to record network activity. B) Normalized power area shown by stimulation frequency (5 Hz or 10 Hz stimulations), frequency band (low theta, 2 – 5 Hz; high-theta, 6 – 12 Hz), and laser wavelength (blue, 473 nm; red, 640 nm). Stimulating interneurons with the blue laser at 5 Hz had no effect on the normalized power area in low-theta, t(3) = 1.03, p = 0.38, and no effect on the normalized power in high-theta, t(3) = -1.07, p = 0.36. Similarly, stimulating interneurons with the blue laser at 10 Hz had no effect on the normalized power area in low-theta, t(3) = 0.69, p = 0.54, nor in high-theta, t(3) = 0.73, p = 0.52. We observed network changes when stimulating interneurons using the red laser. Specifically, when stimulating at 5 Hz, there is a significant difference between normalized power area between laser types in low-theta, t(3.2) = -4.32, p = 0.02, and in high-theta, t(4.1) = -4.14, p = 0.013. When stimulating interneurons at 10 Hz, we saw a significant difference between laser types in high theta, t(4) = -5.76, p = 0.005, but not in low-theta, t(2.4). = -1.39, p = 0.28.

**Supplemental Figure S2.**
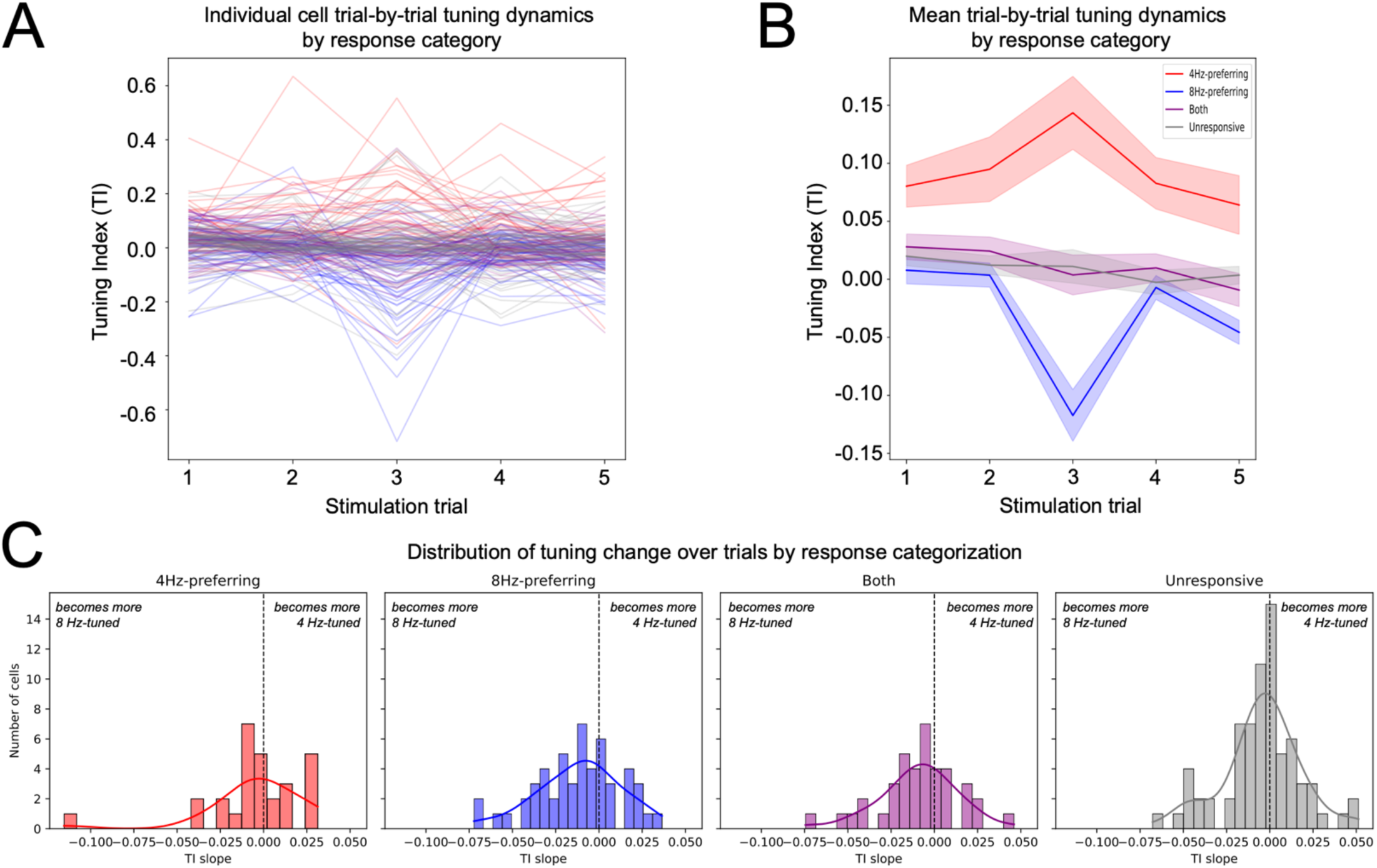
Identified 4 Hz and 8 Hz preferring neurons do not increase or decrease their preference over stimulation trials. A) Tuning Index ([dF/F 4 Hz] – [dF/F 8 Hz]) shown over stimulation trials. Each line represents an individual cell, color coded by eventual preference categorization. B) Mean Tuning Index by response category with SEM shading. C) Histograms and density plots showing distributions of Tuning Index Slope separated by response categorization. Positive slopes represent an increase in 4 Hz tuning while negative slopes represent an increase in 8 Hz tuning. The distributions of identified cells are similarly centered across conditions.

**Supplemental Figure S3.**
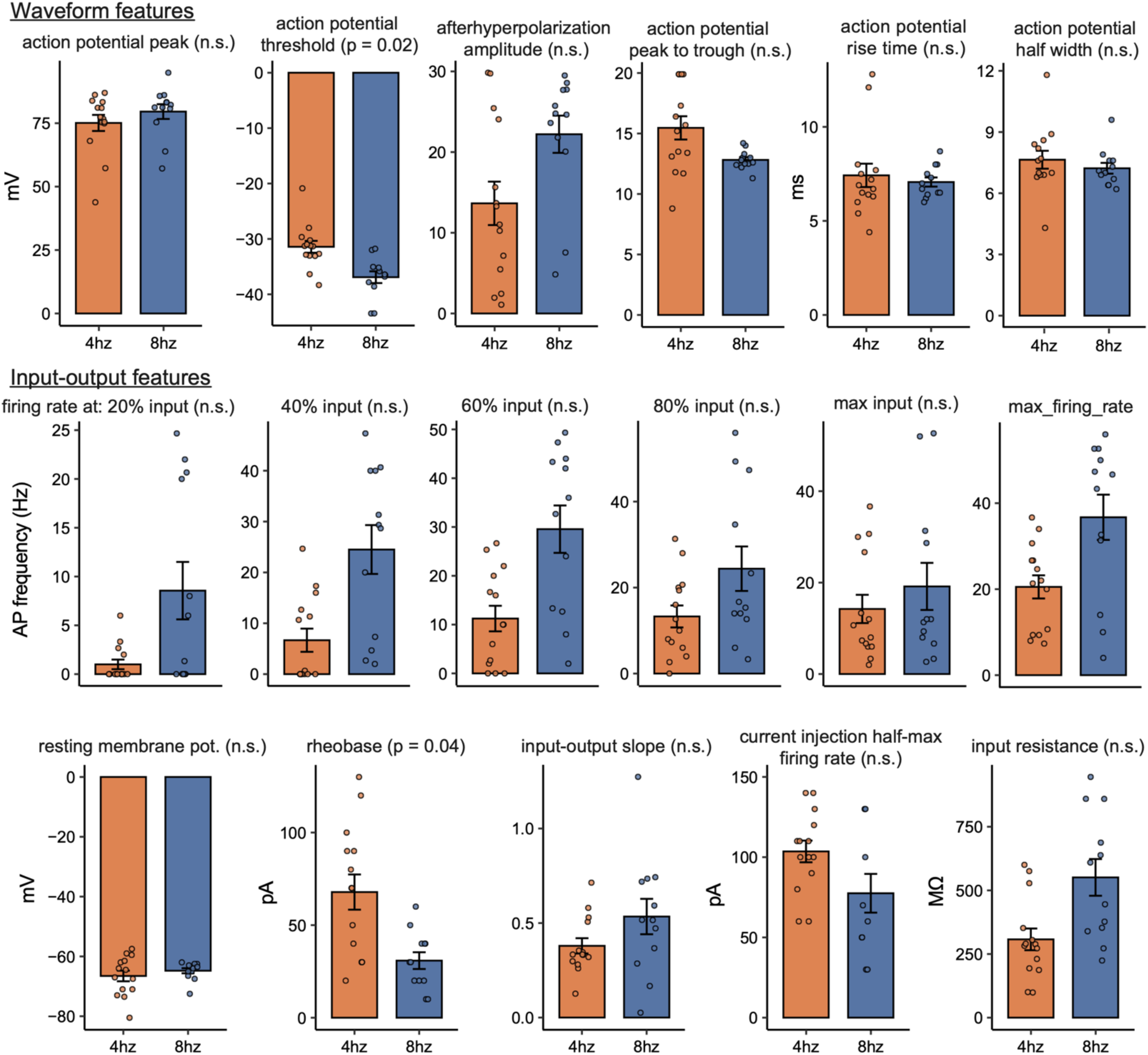
Intrinsic electrophysiological properties of optogenetically-recruited neurons via interneuron stimulations at either 4 Hz (red) or 8 Hz (blue). Top: Waveform properties including, left to right, action potential peak, action potential threshold*, afterhyperpolarization amplitude, action potential peak to trough, action potential rise time, and action potential half-width. Bottom: Input-output features including firing rate at increasing input strengths: 20%, 40%, 60%, 80%, and maximum, as well as the maximum firing rate of recorded neurons. Bottom row, left to right: resting membrane potential, rheobase*, input-output slope, current injection half-max firing rate, and input resistance. * p < 0.05 after Bonferroni corrections for multiple comparisons.

**Supplemental Figure S4.**
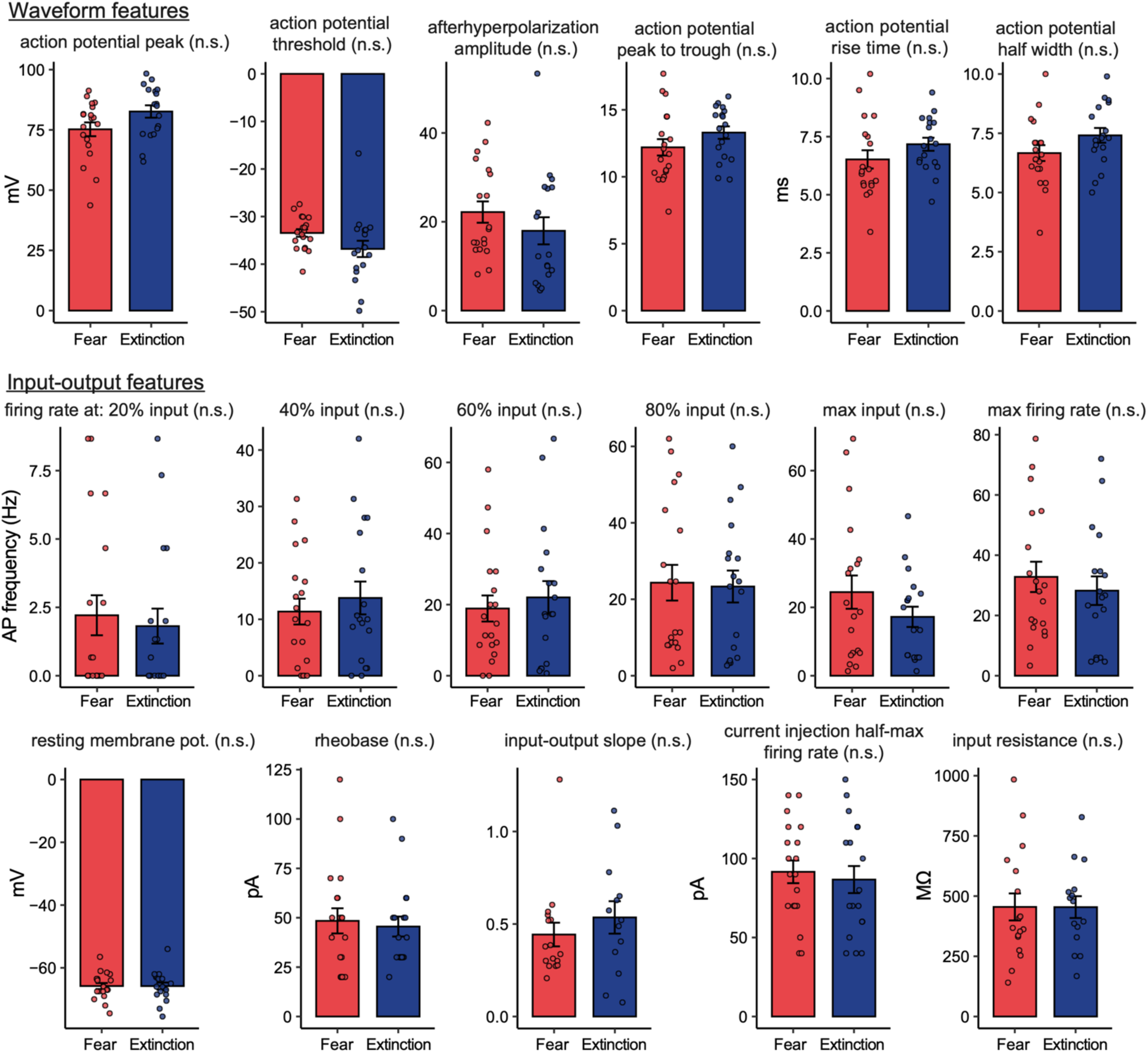
Intrinsic electrophysiological properties of behaviorally-recruited neurons via fear conditioning (red) or extinction learning (blue). Waveform features: Top, Waveform properties including, left to right, action potential peak, action potential threshold, afterhyperpolarization amplitude. Bottom, action potential peak to trough, action potential rise time, and action potential half-width. Input-output features: Top, firing rate at increasing input strengths: 20%, 40%, 60%, 80%, and, Center, maximum, as well as the maximum firing rate of recorded neurons. Current injection half-max firing rate, input resistance, and resting membrane potential. Bottom, maximum firing rate, rheobase, and input-output slope. No significant effects observed.

**Supplemental Figure S5.**
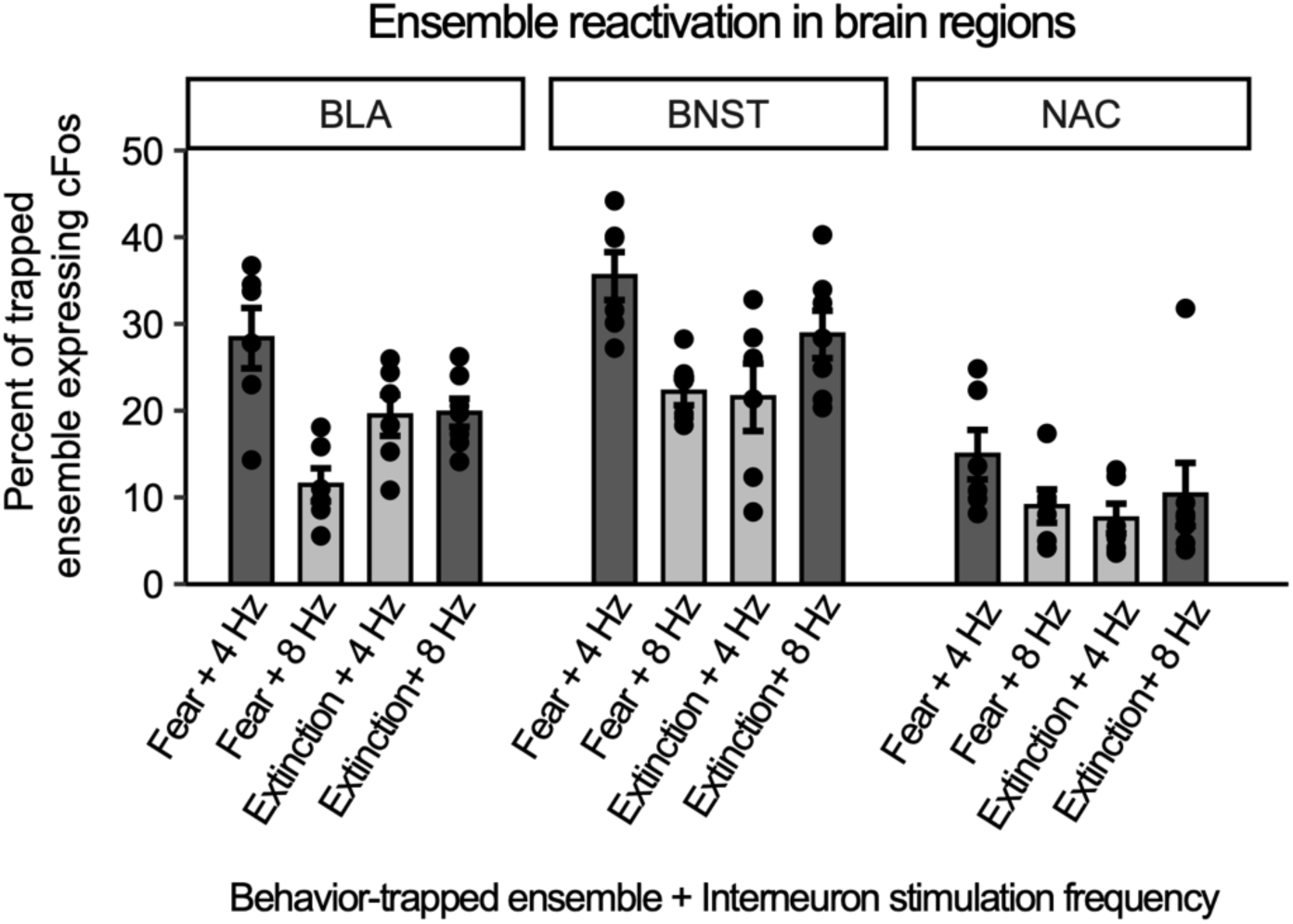
Congruent conditions broken down into behavioral tag method (fear or extinction) and their combination with interneuron frequency stimulation (4 Hz or 8 Hz). Statistical analysis was conducted for data within each brain region, where generalized linear mixed models were used to model ensemble coexpression (coexpression/total_behavioral_ensemble) as a product of behavioral tag and stimulation frequency. For the BLA, there was no significant effect of stimulation frequency on coexpression (OR: 1.19; CI: 0.98 – 1.44; p = 0.085), but there was a significant effect of behavioral tag (OR: 11.04; CI: 1.97 – 61.82; p = 0.006). There was also a significant interaction between stimulation frequency and tag (OR: 0.74; CI: 0.57 – 0.96; p = 0.024). In the BNST, there were no significant effects observed among the behavioral tag (OR: 3.84; CI: 0.59 – 24.94; p = 0.159) nor stimulation frequency (OR: 1.13; CI: 0.92 – 1.39; p = 0.244), nor was there a significant interaction between these variables (OR: 0.88; CI: 0.66 – 1.19; p = 0.412). Similarly, in the NAc, there was not a significant effect of behavioral tag (OR: 6.94; CI: 0.28 – 172.56; p = 0.238) nor stimulation frequency (OR: 1.10; CI: 0.76 – 1.59; p = 0.609), nor was there a significant interaction between these variables (OR: 0.77; CI: 0.47 – 1.28; p = 0.316).

**Supplemental Figure S6.**
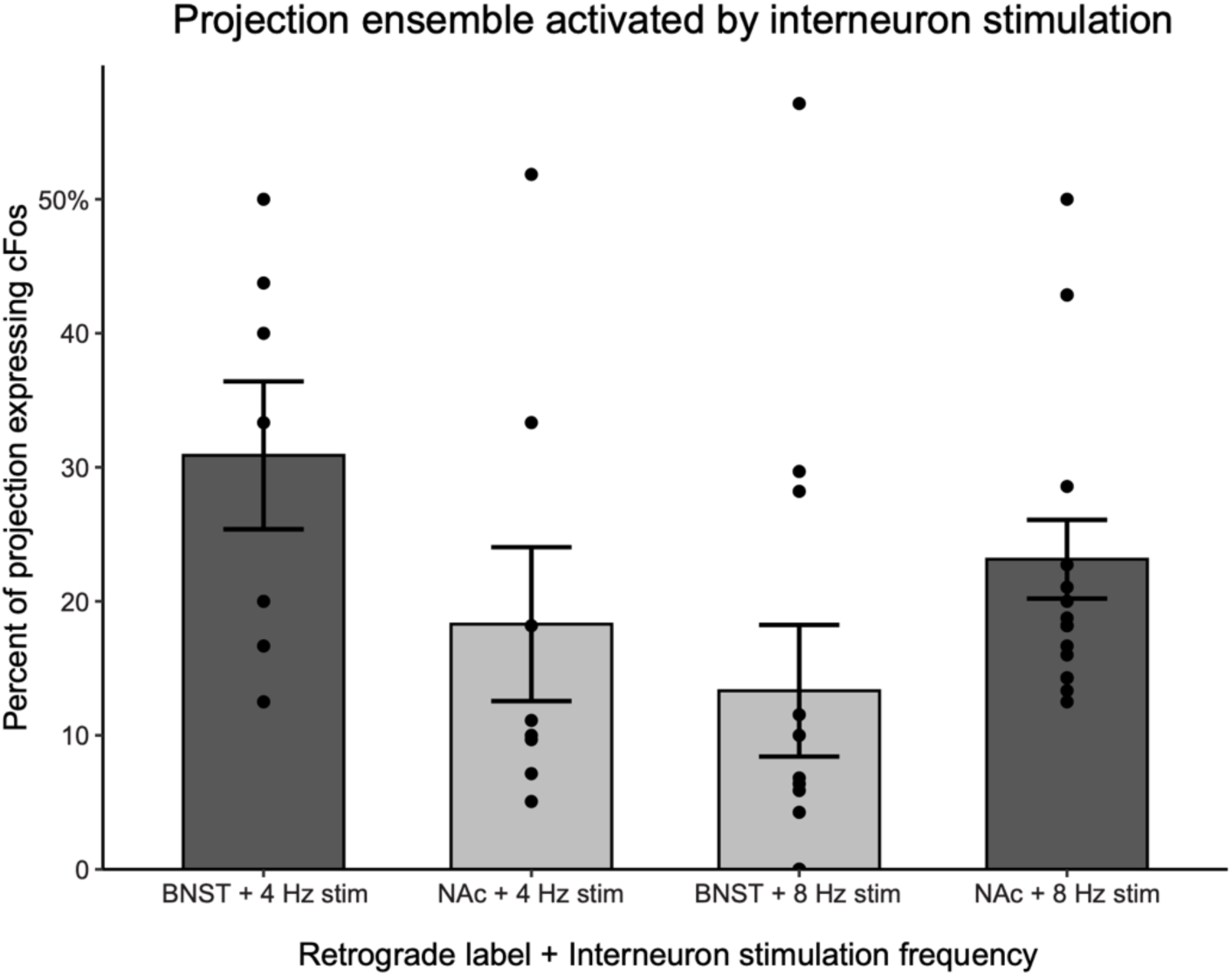
Congruent conditions broken into retrograde label and interneuron stimulation pairings, colored by congruency. Statistical analysis was performed using a generalized linear mixed model weighted by total count of projection neurons observed in the BLA, (cFos coexpression/total) ∼ label * stimulation + (1|animal_id), and family set to binomial. Results showed a significant effect of label on coexpression (OR: 0.11; CI: 0.01 – 0.88; p = 0.037) and a trending label by stimulation interaction (OR: 3.43; CI: 0.99 – 11.88; p = 0.052). There was no effect of interneuron stimulation frequency on coexpression (OR: 0.55; CI: 0.22 – 1.41; p = 0.214).

## Supplemental Methods

### Trial by trial calcium imaging response profiles

To quantify frequency selectivity on a trial-by-trial basis, a tuning index (TI) was computed for each neuron and trial as (R₄Hz − R₈Hz) / (|R₄Hz| + |R₈Hz|), where R₄Hz and R₈Hz represent stimulus-evoked responses. Trial-by-trial changes in tuning were assessed using ordinary least-squares linear regression of TI against trial number for each neuron independently, with the resulting slope parameter used as a measure of within-session response changes. Positive slopes indicate increasing 4 Hz selectivity over time, whereas negative slopes indicate a shift toward 8 Hz selectivity.

For visualization, neurons were sorted according to functional classification and mean stimulus-evoked response magnitude, and ΔF/F responses were organized into per-neuron, per-time-bin matrices separately for each stimulus frequency.

### Intrinsic Properties Analyses

For input-output (IO) curve analysis, only cells that had more than 10 Hz maximum firing rate (5 APs in 0.5 s), were included. The IO slope was calculated using a first-degree least squares polynomial fit (NumPy’s *polyfit*, *n* = 1) between rheobase to 90% of max firing rate. For the IO slope, cells were included only if they fired for at least 3 steps from rheobase to 90% of max APs (at least 3 points for line fit). For waveform analysis, only a maximum of 3 APs were collected from up to the first 3 current steps above rheobase. This was done, to ensure that the afterhyperpolarization (AHP) was maximally preserved. For all analyses, cells were excluded based on their maximum firing rate using Tukey’s outlier detection method (k = 3). To calculate waveform properties, APs were interpolated (SciPy’s *interp1d*, interpolation factor = 10, type = cubic). The AP threshold was detected as 1.5 mV/ms, the AP amplitude was defined as the difference from the AP peak – threshold. The AHP amplitude was defined as the absolute difference between the AHP peak – AP threshold. AP peak-to-trough was calculated as the time between AHP peak to AP peak. The rise time was calculated as the time between AP peak to AP threshold. Statistical tests were conducted in R with Bonferroni corrections for multiple comparisons. Figures were generated using python or R and stylized in Adobe Illustrator.

